# Protocol for Developing a Mouse Model of Post Primary Pulmonary Tuberculosis after Hematogenous Spread in Native Lungs and Lung Implants

**DOI:** 10.1101/2025.02.28.640830

**Authors:** Shivraj M. Yabaji, Suruchi Lata, Igor Gavrish, Ming Lo, Aoife K O’Connell, Hans P Gertje, Colleen E Thurman, Nicholas A. Crossland, Lester Kobzik, Igor Kramnik

## Abstract

This protocol describes a mouse model of post-primary pulmonary tuberculosis (PTB) that develops after hematogenous spread from the primary lesion in native lungs and subcutaneous lung implants. It demonstrates that virulent *Mycobacterium tuberculosis* (Mtb) disseminates to lymphoid tissue in many organs, but selectively damages the lungs. This approach demonstrates a particular vulnerability of the lung tissue to virulent Mtb independent of the route of infection and provides a robust platform for examining lung-specific mechanisms driving TB pathology.

**Highlights:**

1. Mouse model for studying mechanisms driving post-primary pulmonary TB progression in immune hosts
2. Models a hematogenous spread of virulent *Mycobacterium tuberculosis* to the lungs from a primary site of infection
3. Allows for the investigation of lung-specific mechanisms of TB susceptibility using lung tissue implants.

*Graphical abstract:* 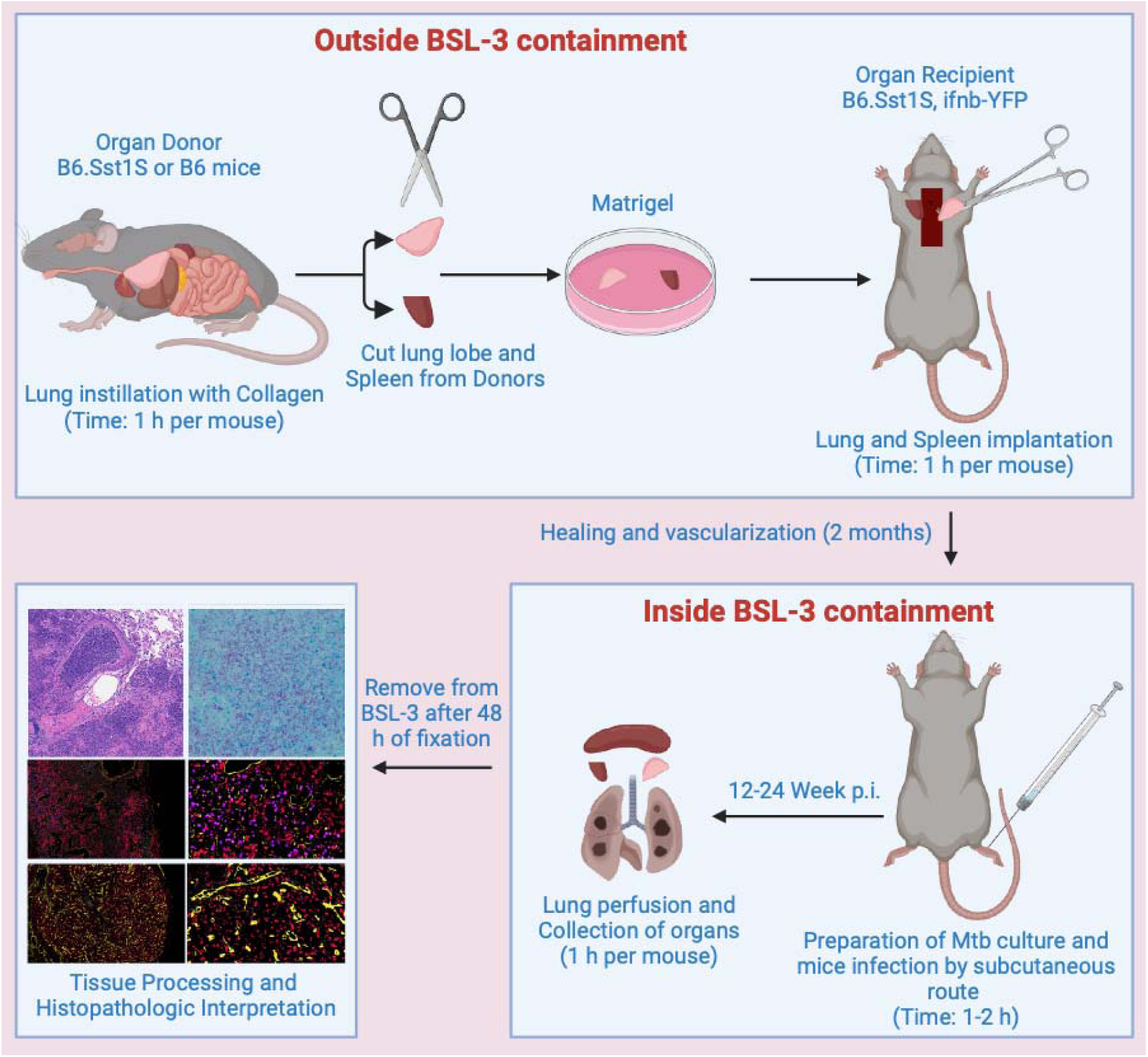

## Before you begin

This protocol describes a mouse model of pulmonary tuberculosis (PTB) using subcutaneous hock infection and lung tissue implantation in B6.Sst1S mice to study hematogenous dissemination, host-pathogen interactions, and genetic factors influencing disease progression.

### Institutional permissions

Obtain institutional permission to perform animal studies and collect tissues under an approved Institutional Animal Care and Use Committee (IACUC) or Institutional Review Board protocol. Our protocol was approved by Boston University’s Institutional Animal Care and Use Committee (IACUC protocol number PROTO201800218).

### Mice

B6J.C3-Sst1C3HeB/Fej Krmn (B6.Sst1S, available from MMRRC stock # 043908-UNC) and B6. Sst1S,ifnb-YFP reporter mice were developed and maintained in our laboratory. C3HeB/FeJ was purchased from Jackson Laboratory (Stock # 000658). C3XB6.Sst1S)F1 was generated in our lab by crossing B6.Sst1S with C3HeB/FeJ.

### Background

Tuberculosis (TB), caused by *Mycobacterium tuberculosis* (Mtb), remains a leading global health concern, causing significant morbidity and mortality. In human patients Mtb spreads hematogenously from primary infection sites and develops post-primary TB lesions in the lungs that lead to severe lung damage and Mtb spread via aerosols. Thus, one of the major challenges in combating TB is understanding mechanisms of TB progression in the lungs.

The development of animal models recapitulating key aspects of human disease is important for studying TB pathogenesis and evaluating novel vaccines and therapies. As in humans, lung is the most vulnerable organ targeted by virulent Mtb in experimental animals. However, modeling pulmonary TB (PTB) after hematogenous spread in immunocompetent hosts was unsuccessful.

In this protocol, we present a mouse model of post-primary PTB using an immunocompetent mouse strain B6.Sst1S mice genetically susceptible to Mtb. In this model mice are infected subcutaneously with virulent Mtb that spread to various organs, but PTB progression occurs specifically in the lungs and lung implants. This model recapitulates pathomorphological features of post-primary PTB lesions in humans, including granulomatous lesions with necrosis and fibronecrotic granulomas. This model allows to study lung-specific mechanisms of PTB progression, including interactions of immune and lung parenchymal cells.

To ensure that pulmonary TB lesions develop after initial priming of adaptive immunity and hematogenous spread, we infected B6.Sst1S mice with virulent Mtb suspension subcutaneously (SQ) in the hock, an established alternative to footpad injection(Kamala, 2007). We have chosen the SQ hock infection to model the hematogenous TB spread, because 1) it allows clear separation of the primary site of infection from secondary, metastatic, lesions; 2) the only anatomically possible route of lung colonization is through hematogenous dissemination, resembling post-primary pulmonary TB progression in immunocompetent humans; 3) SQ hock injection is a common route of immunization(Kamala, 2007), and T cell-mediated immunity is rapidly induced in the regional (popliteal) lymph node after subcutaneous and intradermal immunizations(Apt et al., 1991; Nemeth et al., 2020).

### Mouse Selection and Breeding

1. Use B6.Sst1S mice (B6J.C3-*Sst1^C3HeB/Fej^*Krmn) for this protocol, as this strain is genetically susceptible to TB due to the *susceptible* allele of the *Sst1* locus, which promotes necrotic granulomatous lung inflammation.

2. We maintain a colony of B6.Sst1S mice under specific pathogen-free (SPF) conditions to prevent confounding infections that may impact experimental outcomes.

3. Consider the age of mice, as this can influence the immune response and disease progression. Typically, 8 – 12 week-old mice of both sexes can be used.

### Preparation of Donor Lung or Spleen Tissue for implantation

4. Lung or spleen tissue used for implantation must be obtained from pathogen-free donor mice. Ensure that the genetic background of the donor is appropriately matched for better acceptance of ingrafts.

5. Harvest lung and spleen tissue aseptically to avoid contamination. Use sterile instruments and appropriate biosafety procedures when handling lung tissue to maintain sterility and minimize the risk of external microbial influence.

### Environmental and Biosafety Considerations

6. Perform all procedures involving live Mtb under BSL-3 conditions to ensure researcher safety and prevent environmental contamination.

7. Use personal protective equipment (PPE), including gloves, masks, and lab coats, and follow institutional biosafety protocols for handling infectious materials.

### Animal Handling, Experimental Controls and Reproducibility

8. Ensure appropriate analgesia and post-operative care for mice undergoing lung implantation surgery to reduce pain and distress.

9. Include appropriate controls, such as non-infected mice and mice without lung tissue implants, to account for baseline responses and procedural effects.

10. Standardize experimental conditions, including infection dose, time points for tissue collection, and environmental parameters, to minimize variability.

## Key resources table

| REAGENT or RESOURCE | SOURCE | IDENTIFIER |
| --- | --- | --- |
| **Antibodies** |  |  |
| Rabbit Anti-Ttf1 Monoclonal Antibody, Unconjugated, Clone EP1584Y | Abcam | Cat# ab76013, RRID:AB_1310784 |
| Anti-Uteroglobin antibody (EPR19846) | Abcam | Cat# ab213203, RRID:AB_2650558 |
| Iba1/AIF-1 (E4O4W) XP® Rabbit mAb | Cell Signaling Technology | Cat# 17198, RRID:AB_2820254 |
| **Bacterial and virus strains** |  |  |
| *Mycobacterium tuberculosis* Erdman | ATCC |  |
| Erdman(SSB-GFP, smyc':mCherry) | (Lavin and Tan, 2022) | N/A |
| **Biological samples** |  |  |
| **Chemicals, peptides, antibiotics and recombinant proteins** |  |  |
| DMEM/Ham's F-12 50/50 Mix [+] L-glutamine | Corning® | Cat# 45000-344 |
| Fetal Bovine Serum (FBS) | Hyclone™, | Cat# SH30071.03 |
| Hoechst 33342 | Fisher Scientific | Cat# H3570 |
| Paraformaldehyde Solution 4% in PBS | Fisher Scientific | Cat# |
| L-Glutamine | Corning® | Cat# 25-005-CI |
| Penicillin Streptomycin solution | Corning® | Cat# 30-002-CI |
| HEPES buffer | Corning® | Cat# 25-060-CI |
| L929 Cell Conditioned Media (LCCM) | This paper | N/A |
| Gibco™ 1X DPBS | Fisher scientific | Cat#14-190-235 |
| Middlebrook 7H9 Broth | BD Biosciences | Cat# 271310 |
| Middlebrook 7H10 Agar | BD Biosciences | Cat# 262710 |
| ProLong™ Gold Antifade Mountant | Invitrogen™ | Cat# P36934 |
| Gibco™ Collagen I, rat tail | Fisher Scientific Company | A1048301 |
| Ketamine | Zoetis Inc. | Cat#043-304 |
| Xylazine | Dechra Vet Products, LLC | Cat#047-956 |
| Heparin, sodium salt | Thermo Scientific | Cat#A1619803 |
| 2% Chlorhexidine in 70% Alcohol | GAMA Healthcare Ltd. | Cat#CA2C240 |
| Matrigel Basement Membrane Matrix | Corning® | Cat#354234 |
| Buprenorphine extended-release injectable suspension | Ethiqa XR, Fidelis Animal Health, Inc. | N/A |
| STERIS™ Vesphene™ III se Disinfectant Cleaner | STERIS™ | Cat#647508 |
| 10 mL Control Syringe | BD | Cat#309695 |
| 5 µm filter unit | Millex | Cat#SLSV025LS |
| **Critical commercial assays** |  |  |
| New Fuchsin method | Poly Scientific R&D Corp | Cat#K093 |
| **Deposited data** |  |  |
| **Experimental models: Cell lines** |  |  |
| **Experimental models: Organisms/strains** |  |  |
| Mouse: C57BL/6J | The Jackson Laboratory | Stock No.: 000664 |
| Mouse: B6J.C3-SstI<sup>C3HeB/FeJ</sup>Krmn | (Pichugin et al., 2009) | Stock No: 043908-UNC<br>https://www.mmrrc.org |
| Mouse: C3HeB/FeJ | The Jackson Laboratory | Stock No.: 000658 |
| Mouse: (C3XB6.Sst1S)F1 | This study | N/A |
| Mouse: B6.Sst1S;ifnb-YFP | This study/(Rosenbloom et al., 2022; Scheu et al., 2008) | N/A |
| **Oligonucleotides** |  |  |
| **Recombinant DNA** |  |  |
| **Software and algorithms** |  |  |
| Graphpad Prism 9.5.1 (528) | Graphpad | N/A<br>https://www.graphpad.com/ |
| HALO v3.3.2541.262 | Indica Labs Inc.<br>Clarivate Analytics<br>Oxford Instruments<br>Operatta™<br>Leica VT1200 S | https://indicalab.com/halo/<br>https://endnote.com/downloads<br>[https://imaris.oxinst.com/microscopy-imaging-software-free-trial?](https://imaris.oxinst.com/microscopy-imaging-software-free-trial?source=viewer)<br>source=viewer<br>https://www.perkinelmer.com/in/lab-solutions/product/peretta-cls-system-hh16000020<br>https://www.leicabiosystems.com/us/research/vibratomes/leica-vt1200/ |
| EndnoteX9 | N/A | N/A |
| **Other** |  |  |
| Tissue-Tek VIP-5 automated vacuum infiltration processor | Sakura Finetek | N/A |
| HistoCore Arcadia paraffin embedding machine | Leica | N/A |
| Automate in vivo manual gravity perfusion system for mice double 140 mL – IV 4140 | Braintree Scientific, Inc | Cat# IV 4140 |

### Materials and equipment

#### Media 1

Materials and equipment Media 1
| Reagent | Final concentration | Amount |
| --- | --- | --- |
| DMEM/F12 | - | 430 mL |
| FBS | 10% | 50 mL |
| Penicillin-Streptomycin | 1% | 5 mL |
| 1M HEPES buffer | 20 mM | 10 mL |
| 200 mM L-Glutamine | 2 mM | 5 mL |

To prepare Media 1, add 10 mL of L929 Cell Conditioned Media (LCCM) to every 90 mL above media composition to a final concentration 10%.

#### Collagen I solution

| Reagent | Final concentration | Amount |
| --- | --- | --- |
| Collagen I (3 mg/mL) | 1 mg/mL | 335 μL |
| 10X Media 1 | - | 100 μL |
| 1N NaOH | - | 17 μL |
| PBS | 1X | 548 μL |

**Note-** Prepare freshly and keep it on ice.

**Note-** Prepare freshly and keep it on ice.

#### Ketamine-xylazine solution (1 mL)

| Reagent | Final concentration | Amount |
| --- | --- | --- |
| Ketamine (100 mg/mL) | 10 mg/mL | 100 mL |
| Xylazine (20 mg/mL) | 1 mg/mL | 50 mL |
| PBS | 1X | 0.85 mL |

**Note-** Prepare freshly and keep it at 22-25°C.

**Note-** Prepare freshly and keep it at 22-25°C.

#### Heparin solution

| **Reagent** | **Final concentration** | **Amount** |
| --- | --- | --- |
| Heparin (100 IU/mg) | 10 IU/mL | 50 mg |
| 1X PBS | 1X | 500 mL |

**Note-** Prepare 10X sterile solution and store at 4°C for one month. Prepare 1X solution freshly.

### Equipment

Automate in vivo manual gravity perfusion system for mice double 140 mL – IV 4140 (Braintree Scientific, Inc., Cat# IV 4140) with 20G needles.

## Step-by-step method details

### I. Implant preparation

#### a. Collagen Instillation of Lungs

##### Time: 1h per mouse

1. Anesthetize the mouse using ketamine-xylazine solution according to the weight of the mice (100 μl of ketamine-xylazine solution per 10g) intraperitoneally (i.p) or according to institutional animal care guidelines.

2. Place the mouse on its back and secure it to a Styrofoam board with sterile pins to prevent movement during the procedure.

3. Disinfect the fur on the mouse’s neck and chest with 70% ethanol to ensure a sterile field.

4. Make a midline incision in the skin, starting a few millimeters above the urinary orifice and extending to the chin.

5. Carefully separate the skin from the underlying muscle to expose the trachea.

6. Rotate the Styrofoam board so that the trachea faces the operator.

7. Insert an 18G catheter into the trachea slowly and gently to avoid tissue damage.

8. Remove the metal rod from the catheter once it is in place.

9. Secure the catheter to the trachea using a piece of thread to prevent dislodgment.

10. Load a 5 mL syringe with ice-cold collagen I solution (1 mg/mL).

11. Inflate the lung by slowly injecting the collagen solution through the catheter. Use gentle, even pressure to avoid damaging the lung tissue.

***Critical:*** *A controlled, gradual instillation of Collagen I into the lungs is essential to prevent alveolar rupture while ensuring even distribution*.

12. After inflation, carefully remove the catheter.

13. Tie off the trachea securely to prevent any fluid leakage.

14. Excise the trachea along with the entire inflated lung using sterile dissection tools.

15. Incubate the Lung at room temperature submerged in sterile DMEM F12 media containing 10% FBS and antibiotic.

16. Incubate for 10 minutes to allow the collagen to solidify and stabilize the lung architecture.

17. Trim the lung approximately 10 mm in size using sterile scissors, selecting the left lobe or a suitable right lobe segment.

18. Get the spleen from same mouse and also make the similar size tissue after trimming with sterile scissors.

19. Place the prepared tissue grafts into sterile tissue culture medium containing antibiotics to prevent contamination.

**Note:** Ensure all tools and surfaces are sterile to avoid contamination, and handle the lung tissue gently to preserve its structural integrity.

#### b. Lung and Spleen Implantation

##### Time: 1 h per mouse

20. Anesthetize recipient mice (6 weeks of age or older) with isoflurane and place them in a sternal recumbency position.

21. Prepare the surgical site by shaving the area between the scapulae.

22. Clean the shaved area with 2% chlorhexidine followed by 70% ethanol, repeating the cleaning process three times to ensure sterility.

23. Make a midline incision of approximately 1–2 cm through the skin between the scapulae using sterile surgical tools.

24. Create subcutaneous pockets on both sides of the incision by blunt dissection.

25. Remove the lung and spleen grafts from the antibiotic-containing culture medium.

26. Dip each graft in Corning® Matrigel® Basement Membrane Matrix, LDEV-free (Corning, NY, USA; product number 354234) to coat the tissue.

***Critical:*** *It is important to pre-coat the tissues with Corning® Matrigel® Basement Membrane Matrix before implantation to enhance cell survival, adhesion, and engraftment by providing an extracellular matrix (ECM)-like environment*.

27. Insert the Matrigel-coated lung graft into one subcutaneous pocket.

28. Place the spleen graft (or additional lung graft) into the contralateral subcutaneous pocket.

29. Close the incision using absorbable sutures and secure the skin with skin glue.

30. Administer analgesic treatment with extended-release buprenorphine (Ethiqa XR, Fidelis Pharmaceuticals LLC) subcutaneously at the time of surgery.

31. Allow the mice to recover from anesthesia under close observation.

32. Once recovered, return the mice to their colony housing with their original cage-mates.

33. Monitor the mice post-operatively for any signs of pain or distress, providing additional care as needed.

34. Confirm the acceptance of grafts after 2 months post implantation by manual and visual inspection.

### II. Mtb infection

#### a. Preparation of Mtb for infection

##### Timing-3-4 h

35. The detailed protocol for this step is provided in Yabaji et al., and briefly explained below.

36. Perform all steps in a Biosafety Level 3 (BSL3) facility using approved protocols and disinfectants inside a Biosafety Cabinet (BSC).

37. Dilute 0.5–1.0 mL of frozen Mtb Erdman (SSB-GFP, smyc′::mCherry) stock in 5 mL Middlebrook 7H9 Broth with OADC and transfer to a sterile media bottle.

38. Place the bottle in a secondary container with absorbent paper and a leak-proof lid.

39. Incubate at 37°C in a shaking incubator at 100 rpm for 2–3 days until OD600 is 0.4– 0.5.

40. Transfer the Mtb suspension to a sterile 15 mL conical tube and centrifuge at 2100×g (∼3000 rpm) for 10 minutes at 20°C–25°C.

41. Resuspend the pellet in 5 mL pre-warmed 1X PBS.

42. Sonicate twice for 5 seconds each with a 5–10 second interval in a sonicating water bath.

43. Add 5 mL more pre-warmed 1X PBS, mix, and let sit vertically for 30–60 minutes.

44. Use the upper 8 mL without disturbing the clumps at the bottom.

45. Attach a 5 µm filter to a syringe barrel, place over a conical tube, and transfer the suspension.

46. Insert the plunger and press gently to filter, avoiding gravity filtration to prevent Mtb loss.

47. Measure OD600 of the filtered suspension; expect 0.1–0.2, assuming OD600 = 1 equals ∼3×10^⁸^ bacteria/mL of the single cell suspension of Mtb obtained after filtration.

48. Dilute the Mtb suspension with 1X PBS containing 10^6^ Mtb per 50 μL for mice infection.

49. Plate an aliquot on solid media and count colonies after 3 weeks to confirm the infectious dose.

#### b. Infection of mice

##### Time: 1-2 h

50. Anesthetize the B6.Sst1S mice containing implants as described in step 1.

51. Confirm that the mice are fully anesthetized before proceeding to ensure no movement or discomfort.

52. Place the anesthetized mouse in a restrainer with access to the hind limb.

53. Identify the lateral tarsal region just above the ankle (hock) of the mouse.

54. Using a sterile 28G needle and 1ml syringe, inject 50 μL of the bacterial suspension (prepared in step 48) subcutaneously into the lateral tarsal region of the hind limb.

**Note:** Formation of a bubble is a sign of proper subcutaneous delivery.

55. Monitor the mouse for proper recovery from anesthesia, ensuring it is returned safely to its cage with minimal stress.

### III. Isolation of infected organs for imaging

#### a. Perfusion of lung and collection of organs

##### Time: 30 min per mouse

56. At the desired time points post infection, anesthetize the mice described in step 1.

57. To prevent blood coagulation, inject 200 μl of 1X PBS containing 100 IU/ml heparin intraperitoneally (i.p.).

58. Set up the Automate in vivo manual gravity perfusion system: fill one syringe with 20 ml of 4% PFA in PBS and another syringe with 80 ml of PBS/heparin solution (10 U/ml). Attach a 20G needle.

59. Pin the mouse securely to a Styrofoam board to immobilize it.

60. Disinfect the fur thoroughly with 70% ethanol using a squeeze bottle.

61. Make a midline incision from just above the urinary orifice up to the chin.

62. Separate the skin from the underlying muscle layer to expose the abdominal area and cut peritoneum to expose intestine. Do not open the rib cage to prevent lung collapse.

63. Gently move the intestines to the left side to expose the inferior vena cava, ensuring no organs obstruct blood flow during perfusion.

64. Cut the skin near the lower limbs to facilitate blood drainage, maintaining continuous flow to prevent coagulation (avoid a “blood swamp”).

65. Position the dissection board upright within the tray.

66. Make small incisions to cut the inferior vena cava and aorta in abdominal area.

67. Rapidly insert a 20G needle attached to the Automate in vivo manual gravity perfusion system into the retro-orbital sinus, adjusting as needed, and begin perfusion with PBS + Heparin using the perfusion assembly.

68. Observe pulse-like blood flow, perfuse until blood is replaced by clear PBS. Use approximately 20 mL of PBS + Heparin mix.

69. Switch to 4% PFA and continue perfusing with about 10 mL of fixative.

**Note:** Before starting the procedure of next mouse flush the system with 5 mL of PBS + Heparin to remove any PFA residues.

70. Lower the dissection board and place tools in a Falcon tube filled with 1% Vesphene.

71. Hold the sternum with tweezers and make a small hole in the diaphragm to collapse the lungs. Carefully cut the rib cage, do not to cut the lung.

72. Extend cuts along the diaphragm to fully open the thoracic cavity. Make a midline incision from the sternum to the chin, spread the rib cage, and pin it open, avoiding sharp parts.

73. Rotate the board so the trachea faces the operator. Place a thread beneath the trachea.

74. Slowly insert an 18G catheter into the trachea without force, remove the metal rod, and place it in 1% Vesphene. Tie the catheter in place.

75. Attach an insulin syringe with 1 mL of PBS and inflate the lungs. Collect bronchoalveolar lavage (BAL) fluid if required. Repeat 3 - 5 times to remove residual air. Do not overinflate and avoid lung rupture.

76. Inflate the lungs slowly with 2mL of PBS. Remove the catheter and tie off the trachea immediately.

77. Isolate the lungs and place them in a 50 mL Falcon tube with 10% formalin. Flip the tube upside down to ensure complete submersion.

78. Collect other organs (spleen, liver, kidney, gut, lymph nodes, and any required for study) and place them in 50 ml Falcon tube filled with 10% formalin and palce the tube.

79. Remove pins, reverse the mouse to face down, and pin the limbs. Open the back skin with scissors and forceps to collect implanted tissues and place them in 10% formalin.

80. Place all dissection tools in 1% Vesphene solution and clean the biosafety cabinet.

81. Fix the tissues for 24 hours, remove from BSL-3 containment and transfer in sterile 1X PBS in 50 ml Falcon tube.

#### b. Tissue Processing, and Histopathologic Interpretation

82. Perform the tissue processing and histopathological interpretation using standard classical methods.

83. Stain a subset of tissue section for acid fast bacteria (AFB).

84. For brightfield acid-fast staining of Mtb-infected lung sections, use the New Fuchsin method (Poly Scientific R&D Corp., Cat. No. K093, Bay Shore, NY, USA) and follow the manufacturer’s instructions for the staining procedure.

**Alternatives:** You can use other methods of Mtb staining like fluorescent Acid-Fast Bacteria (AFB) Staining using Auramine-Rhodamine Method.

85. If required the lungs can be processed for 3D imaging using thick sections (Lata et al., 2025).

## Expected outcomes

### I. Progression of PTB in Native lungs of B6.Sst1S

In pilot studies, we infected adult female mice via subcutaneous hock injection with 10^4^, 10^5^ and 10^6^ CFU of Mtb Erdman and only 10^6^ CFU induced pulmonary granulomas in 100% of animals by 12 weeks post-infection (wpi) (**Table 1**). However, when we infected the wild type B6 and congenic B6.Sst1S mice with 10^6^ CFU and observed the weight loss and decreased survival only in the B6.Sst1S mice (Yabaji et al., 2023). Groups of B6 and surviving B6.Sst1S mice were sacrificed at 11- and 20-weeks post-infection (wpi), developing PTB lesions in both strains. Based on Mtb load and histopathological features in the pulmonary lesions we categorized the lesions in two types, classified as “paucibacillary” and “multibacillary”. Majority of B6.Sst1S develop multibacillary pulmonary lesions while B6 mice only develop paucibacillary pulmonary lesions (**Table 2**). B6 mice develops only minimal to mild granulomatous interstitial lesions containing rare single acid-fast bacilli-AFB, while B6.Sst1S lesions displayed a broad variability of lesions including severe lesions characterized by the presence of cholesterol clefts, neutrophils and areas of necrosis with innumerable AFB (Yabaji et al., 2023) (**Fig. 1**). Moreover, we compared the males and females B6.Sst1S mice for Mtb infection and similar disease progression was observed in both sexes (**Table 3**). These findings emphasize the role of *Sst1* locus in determining pulmonary TB severity and survival.

**Figure 1:**
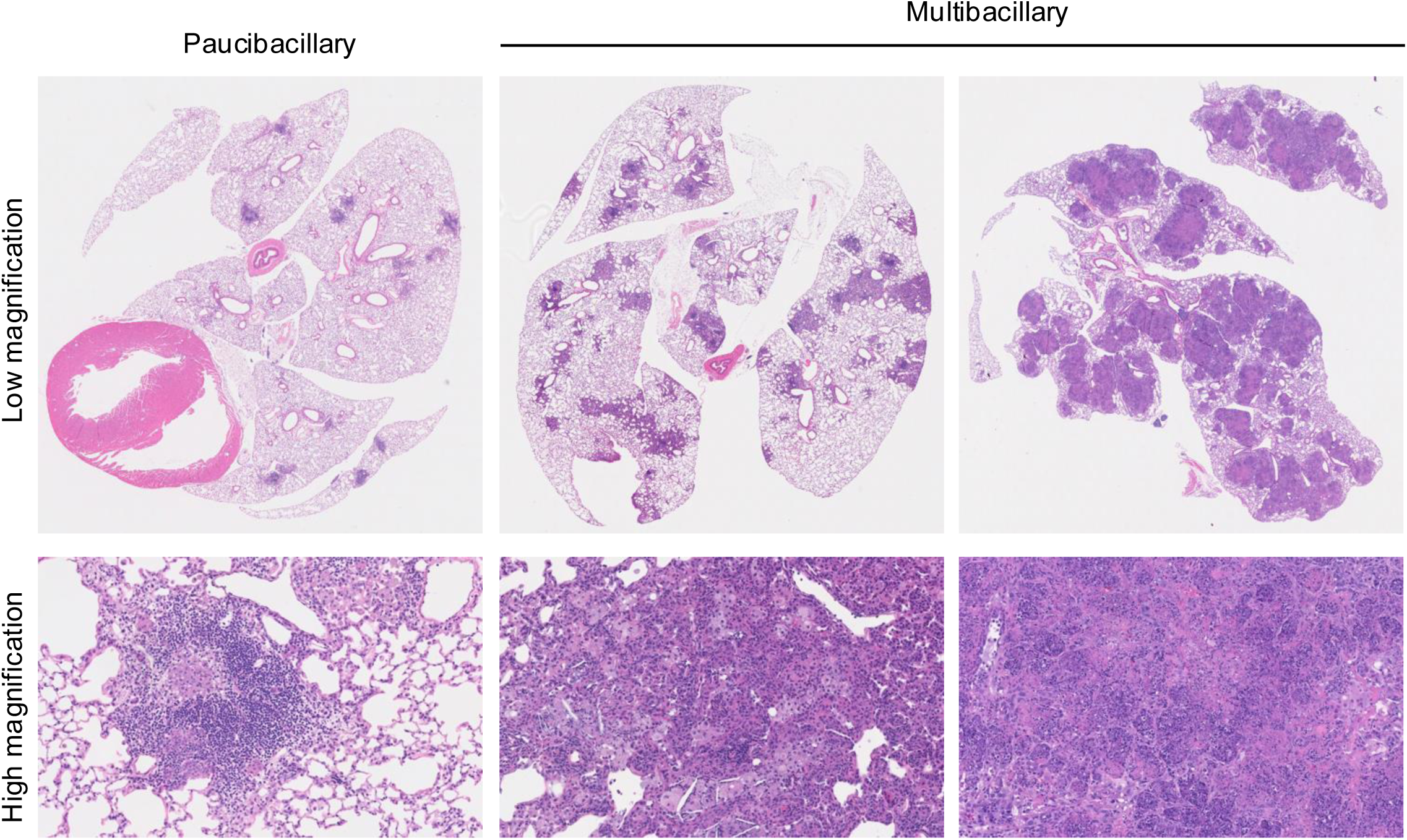
Progression of PTB lesions in Native lungs of B6.Sst1S. Stages of PTB progression in B6.Sst1S mice following subcutaneous Mtb infection. Representative low-magnification (1X) and corresponding high magnification histopathology images of lung sections (H&E staining).

**Table 1.**
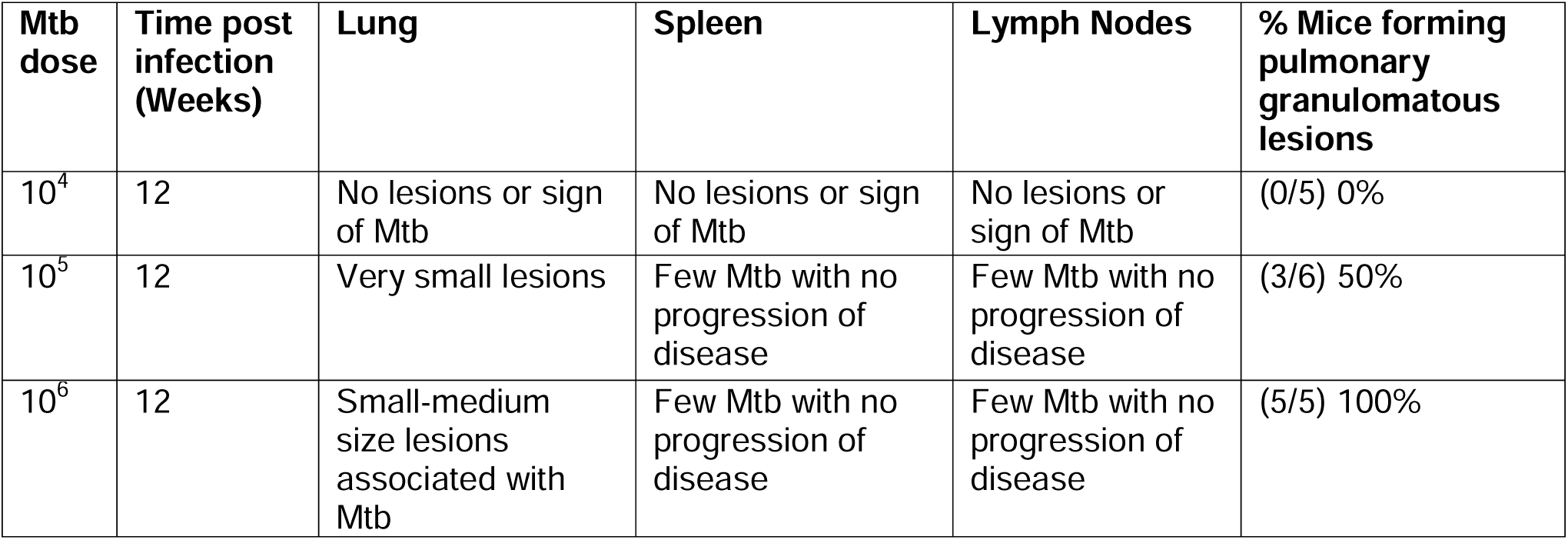
Dose-dependent progression of Mtb infection in B6.Sst1S mice.

**Table 2.** Effect of genetic background on pulmonary TB lesions progression in mice within 11-20 weeks post infection.

| **Genotypes** | **Sex** | **Multibacillary** | **Paucibacillary** | **No lesions** | **Total** |
| --- | --- | --- | --- | --- | --- |
| B6 | Female | 0 (0%) | 7 (77.7%) | 2 (22.23%) | 9 |
| B6.Sst1S | Female | 24 (92%) | 2 (8%) | 0 (0%) | 26 |

**Table 3.** Effect of sex on pulmonary TB lesions progression.

| **Genotypes** | **Sex** | **Multibacillary** | **Paucibacillary** | **No lesions** | **Total** |
| --- | --- | --- | --- | --- | --- |
| B6.Sst1S | Male | 6 (75%) | 2 (25%) | 0 | 8 |
| B6.Sst1S | Female | 9 (64.3%) | 3 (21.4%) | 2 (14.3%) | 14 |

### II. Effect of Genetic Background and age on TB Progression in native lungs

We further investigated the genetic influence on tuberculosis (TB) progression in different sst1-susceptible mice, using B6.Sst1S, C3HeB/FeJ (HeB), and F1 hybrids (HeB x B6.Sst1S). We infected all strains of mice (aged 10-12 week) with Mtb at 10^6^ CFU subcutaneously. By 12–20 all strains developed TB lesions; however, B6.Sst1S mice displayed more multibacillary lesions with necrosis compared to HeB and F1 hybrids (**Table 4**). We also observed survival differences and B6.Sst1S mice showed 67% survival with weight loss, whereas HeB and F1 hybrids demonstrated 100% survival rates and no significant weight loss throughout the course of infection (**Fig. 2A - B**). We also measured Mtb loads in several organs and observed the highest bacterial burden in the lungs of B6.Sst1S mice, with bacterial dissemination to secondary organs, including the spleen, liver, lymph nodes, and gut (**Fig. 2C**). In the extrapulmonary organs Mtb is rarely present and localized exclusively within small macrophage aggregates. Additionally, to confirm the effect of age on disease progression, we selected the F1 hybrid strain with the least disease severity and infected young and adult (HeB × B6.Sst1S)F1 mice with Mtb via the subcutaneous route using 10^6^ CFU. Interestingly, younger F1 mice exhibited more multibacillary lesions compared to their adult counterpart (**Table 5**). These findings emphasize the critical role of genetic background in determining pulmonary TB severity and survival. Host genetics significantly impact lesion control, bacterial load, and disease progression.

**Figure 2.**
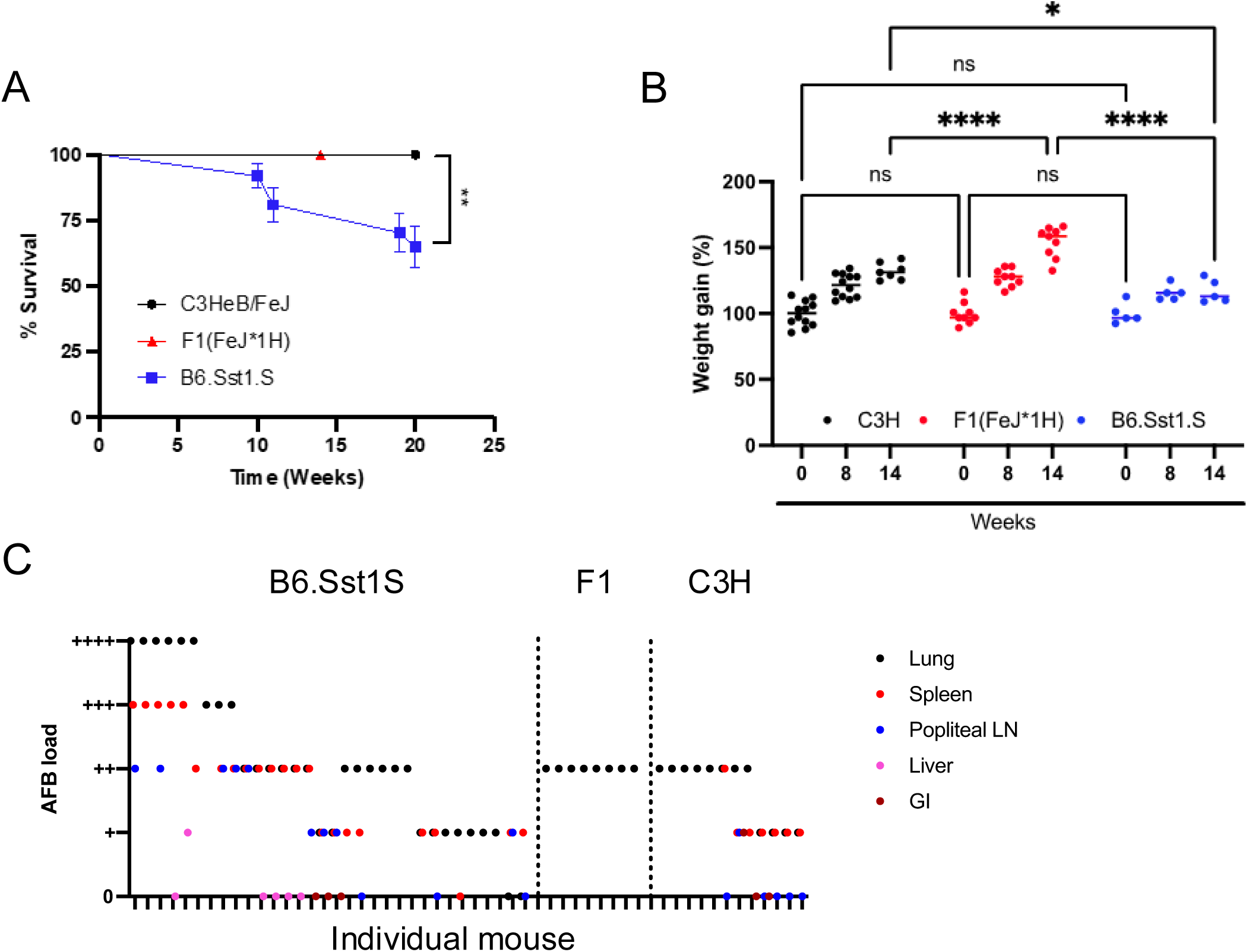
Comparison of post-primary pulmonary TB progression in *sst1* susceptible B6.Sst1S, C3HeB/FeJ mice and their F1 hybrids. **A.** Survival curves of mice infected with M. tuberculosis. Survival of C3HeB/FeJ (n=12), F1 (FeJ*1H) (n=23) and B6.Sst1S (n=37) following hock infection with 10^6^ CFU of M. tuberculosis Erdman(SSB-GFP, smyc′::mCherry). The Log-rank (Mantel-Cox) test was applied to determine the statistical significance. **B.** Weight of the infected mice: C3HeB/FeJ (n=12), F1 (FeJ*1H) (n=23) and B6.Sst1S (n=37) following hock infection with 10^6^ CFU of *M. tuberculosis* Erdman(SSB-GFP, smyc′::mCherry). The statistical significance was performed by two-way ANOVA using Tukey’s multiple comparison test. **C.** Semi-quantitative acid-fast bacteria (Mtb) loads in the lungs, spleens , popliteal lymph nodes and livers of the infected mice at 14 weeks post infection. Each hash on the X-axis represents individual mouse. The p value ≤0.05 was considered statistically significant. Significant differences are indicated with asterisks (*, P < 0.05; **, P < 0.01; ***, P < 0.001; ****, P < 0.0001).

**Table 4.**
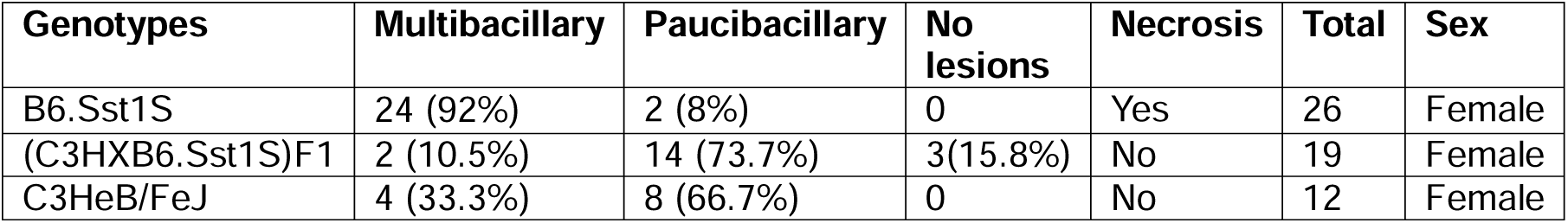
Effect of genetic background on pulmonary TB lesions progression in *sst1* susceptible mice within 12-20 weeks post infection.

**Table 5.** Effect of age on pulmonary TB lesions progression in *sst1* susceptible mice within 12-20 weeks post infection.

| **Genotypes** | **Age** | **Multibacillary** | **Paucibacillary** | **No lesions** | **Total** | **Sex** |
| --- | --- | --- | --- | --- | --- | --- |
| (C3HXB6.Sst1S)F1 | Adult | 2 (10.5%) | 14 (73.7%) | 3(15.8%) | 19 | Female |
| (C3HXB6.Sst1S)F1 | Young | 4 (57.1%) | 2 (28.6%) | 1(14.3%) | 7 | Female |

### III. Investigating TB progression in lung and spleen implants

We investigated the progression of PTB lesions in the native lungs after hematogenous spread under pre-existing condition. Further we wanted to study lung-specific TB progression using subcutaneous implantation of lung tissue fragments from B6.Sst1S into B6.Sst1S mice to study the effect of oxygenation and lung microenvironment. We co-implanted spleen fragments as controls. Additionally, we compared the lung implants with 1% collagen instillation or without instillation before implantation. Two months post-implantation, mice were infected via the subcutaneous hock route, and tissues were collected, and we observed a better preservation of structural integrity of tissue in collagen instilled condition (**Fig. 3**).

**Figure 3:**
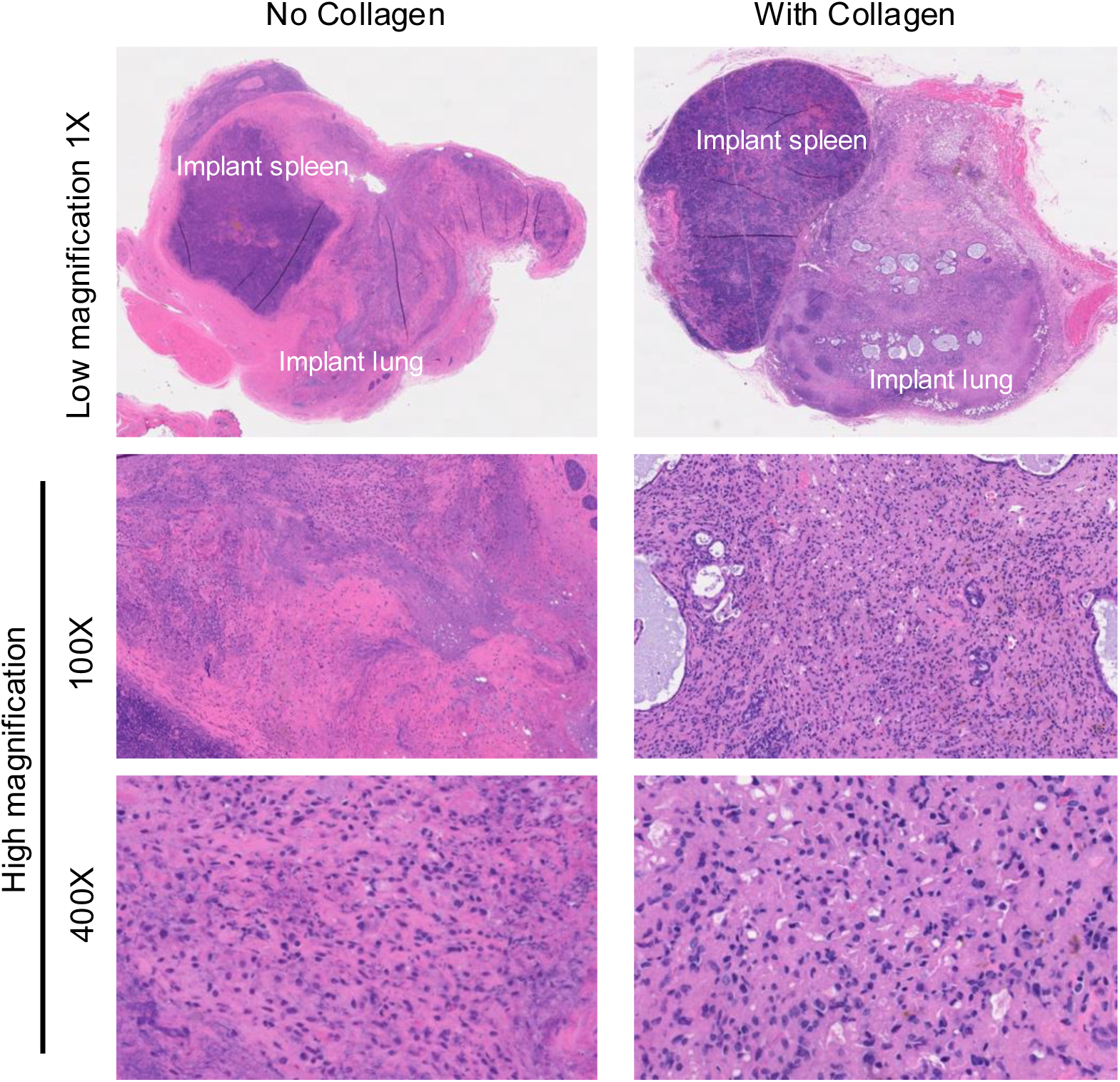
Effect of Collagen I instillation on lung implant condition. H&E-stained images at low and high magnification of implanted lungs from B6.Sst1S mice with or without collagen instillation. 1% Collagen I solution in tissue culture medium was instilled prior to implantation into recipient mice. B6.Sst1S mice were hock infected with 10^6^ CFU of Mtb and sacrificed at 20 wpi.

We observed the formation of organized necrotic granulomas and fibrotic capsules with innumerable Mtb in implanted lung, while spleen implants rarely show bacteria without granulomas (**Fig. 4A - C**). Using fmIHC, we confirmed the presence of lung specific markers in the lung implants including epithelial markers Nkx2.1 (Ttf1), uteroglobin and we also included macrophage marker, Iba1. The lung specific markers were exclusively present in the implanted lung but not in the implanted spleen. However, Iba1 was expressed in both implanted tissues (**Fig. 4D**).

**Figure 4.**
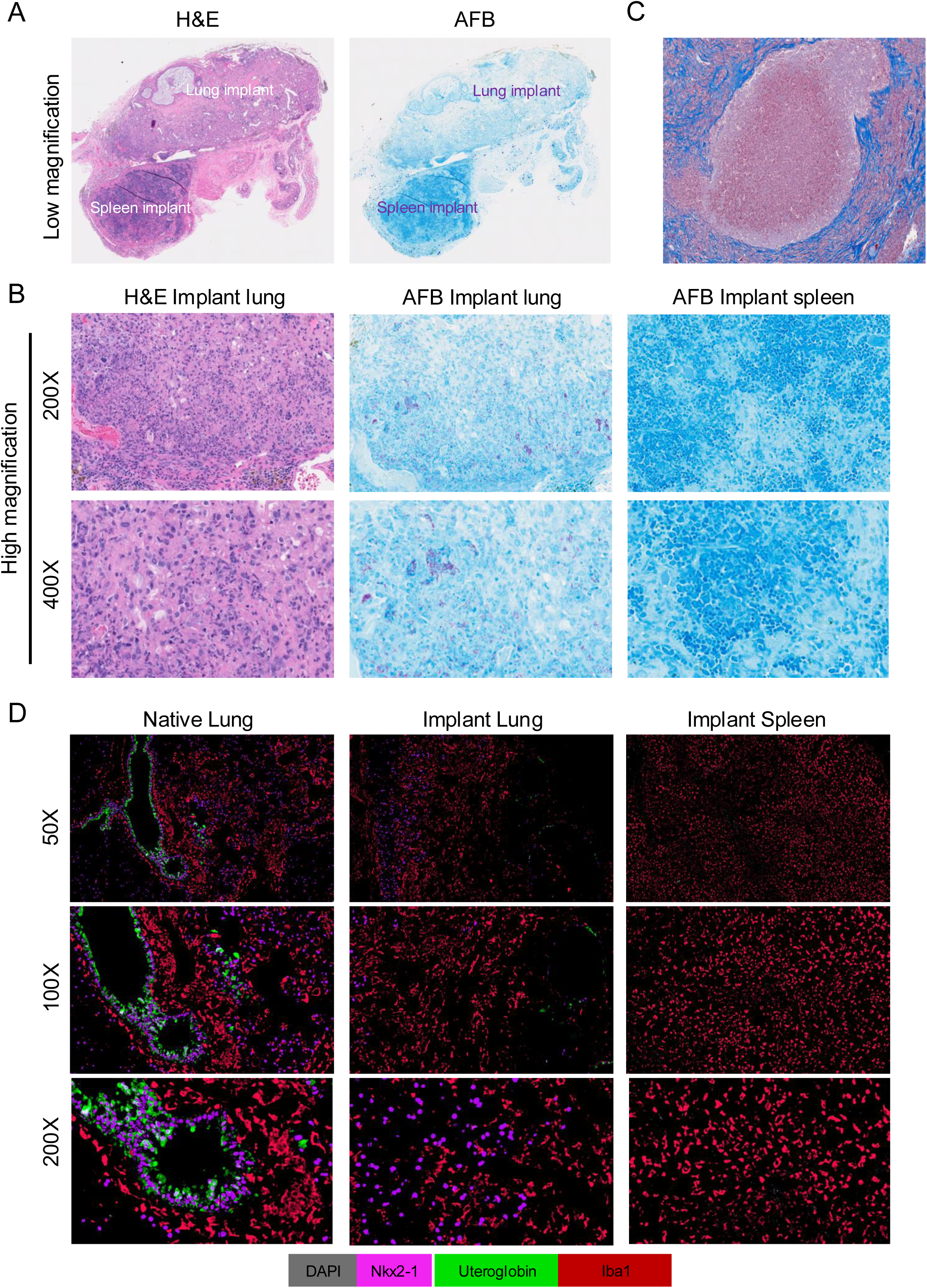
TB lesions in lung implants. **A** and **B**. Low and high magnification images showing H& E and AFB staining of implanted lung and spleen. The Mtb and Mtb induced lesions were only present in implanted lung but not in implanted spleen. **C**. Masson’s Trichrome staining of implanted lung showing fibrotic capsule surrounding necrotic granuloma in a lung implant. 100X magnification. **D**. Multiplex fluorescent immunohistochemistry (mfIHC) of native lung vs lung and spleen implants of Mtb infected B6.Sst1S mice with lung epithelial cell markers Ttf1 (Nkx2.1) and Uteroglobin and a macrophage marker Iba1.

We also observed a correlation between the native and implanted lungs. Paucibacillary lesions in the native lung were associated with an absence of lesions and Mtb in the implanted lungs. In contrast, multibacillary lesions in the native lungs led to lesion formation and Mtb replication in the implanted lungs (**Fig. 5A - B**).

**Figure 5.**
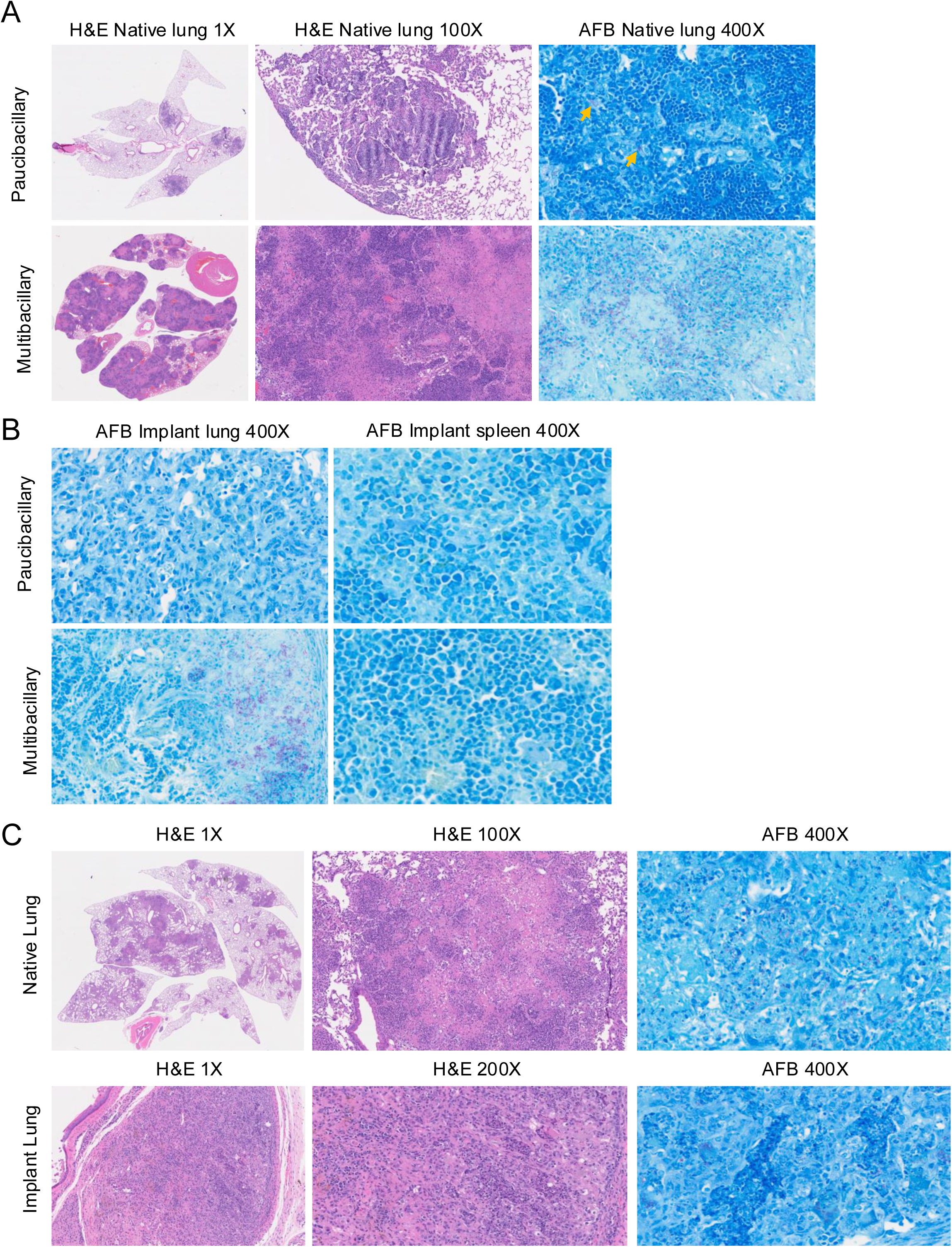
Correlation between disease progression in native lungs and implanted lungs. **A.** H&E and AFB images of pauci- and multibacillary TB lesions in native lungs **B.** AFB staining of the corresponding lung and spleen implants isolated from the same mice presented in A. Multibacillary TB lesions were found only in the lung implants of mice with multibacillary lesions in their native lungs. **C.** H&E and AFB images showing TB lesions in the native lung of B6.Sst1S recipient mice (upper panel) and in their implanted lung tissue isolated from B6 donors mice (lower panel). Implanted B6 lung tissue developed the necrotic lesions containing numerous Mtb .

To investigate the effect of the recipient mice’s genetic background on implanted lung tissue from wild-type B6 mice, we subcutaneously implanted lung tissue from wild-type B6 mice into B6.Sst1S mice. After recovery, these mice were infected with 10^6^ CFU of Mtb Erdman. At 20 weeks post-infection, the implanted B6 lung also developed necrotic lesions with numerous Mtb (**Fig. 5C**).

## Quantification and statistical analysis

To compare multiple groups with two or more variables, we conducted a two-way analysis of variance (ANOVA) and applied adjustments for multiple post hoc comparisons. Various comparisons were made, including comparisons among all groups, between the control group and all other groups, and between selected groups. For comparisons involving multiple groups with a single variable, a one-way ANOVA was used, with corrections for multiple post hoc comparisons. In cases where only two groups were compared, two-tailed paired or unpaired t-tests were employed. The Log-rank (Mantel-Cox) test was utilized to assess statistical significance in mouse survival data. All statistical analyses were performed using GraphPad Prism 9 software. A p-value of < 0.05 was considered statistically significant, and statistical significance was indicated by asterisks (*, P < 0.05; **, P < 0.01; ***, P < 0.001; ****, P < 0.0001).

## Limitations

The subcutaneous infection model does not recapitulate the natural route of infection in the majority of humans. However, it allows for clear separation between the primary and post-primary “metastatic “ pulmonary TB lesions and demonstrates lung vulnerability to Mtb irrespective of the route of infection. The lung implantation model complements this by enabling the study of the lung tissue-specific elements promoting TB progression, although it cannot fully replicate the native lung’s architecture, vascularization, oxygenation, and ventilation potentially impacting TB progression. Implant stability is influenced by tissue size and preparation, as smaller fragments are often resorbed. Histocompatibility of the host and lung implant need to be considered.

## Troubleshooting

### Problem 1

Appropriate dose of Mtb to establish a reliable and reproducible subcutaneous infection in mice, ensuring consistent infection dynamics while minimizing variability and overexposure.

### Potential solution

Use the optimum dose (10^6^) of *Mtb* Erdman to consistently induce pulmonary granulomatous lesions in 100% of mice by 12 weeks post-infection (wpi) (**Table 1**).

### Problem 2

Selection of mice background to get pulmonary necrotic granulomatous lesions by subcutaneous infection.

### Potential solution

Use B6.Sst1S mice (B6J.C3-*Sst1^C3HeB/Fej^*Krmn) to induce pulmonary necrotic granulomatous lesions through subcutaneous infection in the hock. Monitor lesion progression, which drives Mtb replication. Use of B6, C3H, or F1 mice results into formation of pulmonary lesions with reduced bacterial loads and tissue damage.; however, these lesions do not progress to necrotic granulomatous lesions (**Table 4**).

### Problem 3

Disappearance or collapse of smaller implanted tissues from recipient mice.

### Potential solution

Observed that smaller tissue sections disappear during recovery and vascularization, failing to establish stable implants. Use about half of the left lobe or one of the right lobes to implant which remain stable and persist throughout recovery and critically determines the success of lung implantation.

### Problem 4

How can structural integrity of lung tissue be maintained during implantation to improve implantation success?

### Potential solution

Instill lungs with 1% collagen before implantation to support and preserve tissue structure. Collagen-instilled lung implants show significantly better structural integrity compared to untreated controls, demonstrating that collagen instillation is essential for maintaining lung architecture and enhancing implantation outcomes (**Fig. 3**).

### Problem 5

Variation in PTB progression in individual mice after hock infection.

### Potential solution

To detect organized granulomas in lung implants, monitor weight and clinical scores weekly and sacrifice mice when they start losing weight, typically around 12–24 weeks post-infection. This timing increases the likelihood of obtaining lung implants with lesions and detectable Mtb (**Fig. 5A**). Implant spleen or other non-lung tissues to verify the specificity of lung susceptibility to TB infection following hematogenous spread.

## Resource availability

### Lead contact

Further information and requests for resources and reagents should be directed to and will be fulfilled by the lead contact, Igor Kramnik.

### Technical contact

Technical questions on executing this protocol should be directed to and will be answered by Shivraj M. Yabaji

## Materials availability

All unique/stable reagents generated in this study are available from the Lead Contact with a completed Materials Transfer Agreement.

## Data and code availability

This study did not generate datasets/code and any additional information will be available from the lead contact upon request.

## Acknowledgments

The work was supported by NIH R01HL126066 (I.K.) and NIH grants S10OD026983 and S10OD030269 (N.A.C.).

## Author contributions

Conceptualization, IK, SMY; methodology, SMY, ML, SL, IG, AOC, HPG, CET; validation and formal analysis, SMY, ML, NAC; investigation, SMY, SL, IG, LK, NC; resources, NC, IK; writing – original draft, SMY; writing – review & editing, IK, LK, NAC; supervision, IK; funding acquisition, IK, NAC and LK

